# Prevalence of near-death experiences and REM sleep intrusion in 1034 adults from 35 countries

**DOI:** 10.1101/532341

**Authors:** Daniel Kondziella, Markus Harboe Olsen

## Abstract

**Background:** Near-death experiences have fascinated humans for centuries, but their origin and prevalence remain unknown.

**Methods:** Using an online crowdsourcing platform, we recruited 1034 lay people from 35 countries to investigate the prevalence of near-death experiences and self-reported REM sleep intrusion. Reports were validated using the Greyson Near-Death Experiences Scale (GNDES) with a score of ≥7 as cut-off point for identifying near-death experiences.

**Results:** Near-death experiences were reported by 106 of 1034 participants (10%; CI 95% 8.5-12%). REM sleep intrusion was more common in people with near-death experiences (n=50/106; 47%) than in people with experiences with 6 points or less on the GNDES (n=47/183; 26%) or in those without any such experience (n=107/744; 14%; p=<0.0001). Following multivariate regression analysis to adjust for age, gender, place of origin, employment status and perceived danger, this association remained highly significant; people with REM sleep intrusion were more likely to exhibit near-death experiences than those without REM sleep abnormalities (odds ratio 2.85; CI 95% 1.68-4.88; p=0.0001).

**Conclusions:** The prevalence of near-death experiences in the public is around 10%. While age, gender, place of residence, employment status and perceived threat do not seem to influence the prevalence of near-death experiences, there is a significant association with REM sleep intrusion. This finding is in line with the view that despite imminent threat to life, brain physiology must be well-preserved to perceive these fascinating experiences and store them as long-term memories.

## INTRODUCTION

Near-death experiences can be defined as extraordinary conscious perceptual experiences, including emotional, self-related, spiritual and mystical experiences, occurring in a person close to death or in situations of imminent physical or emotional threat.^1^ Reports of near-death experiences include, but are not limited to, increased speed of thoughts, distortion of time perception, out-of-body experiences, and visual and auditive hallucinations. ^1–3^

Although near-death experiences have fascinated humans for centuries, it remains unknown how prevalent they are and what causes them. However, these experiences share phenotypical features with those made during rapid eye movement (REM) sleep (**Table 1**).^4^ Here, we recruited a large multinational sample of lay people from a crowdsourcing platform to estimate the prevalence of near-death experiences and to test the hypothesis that these experiences are associated with a propensity for REM sleep intrusion.

**Table 1.**
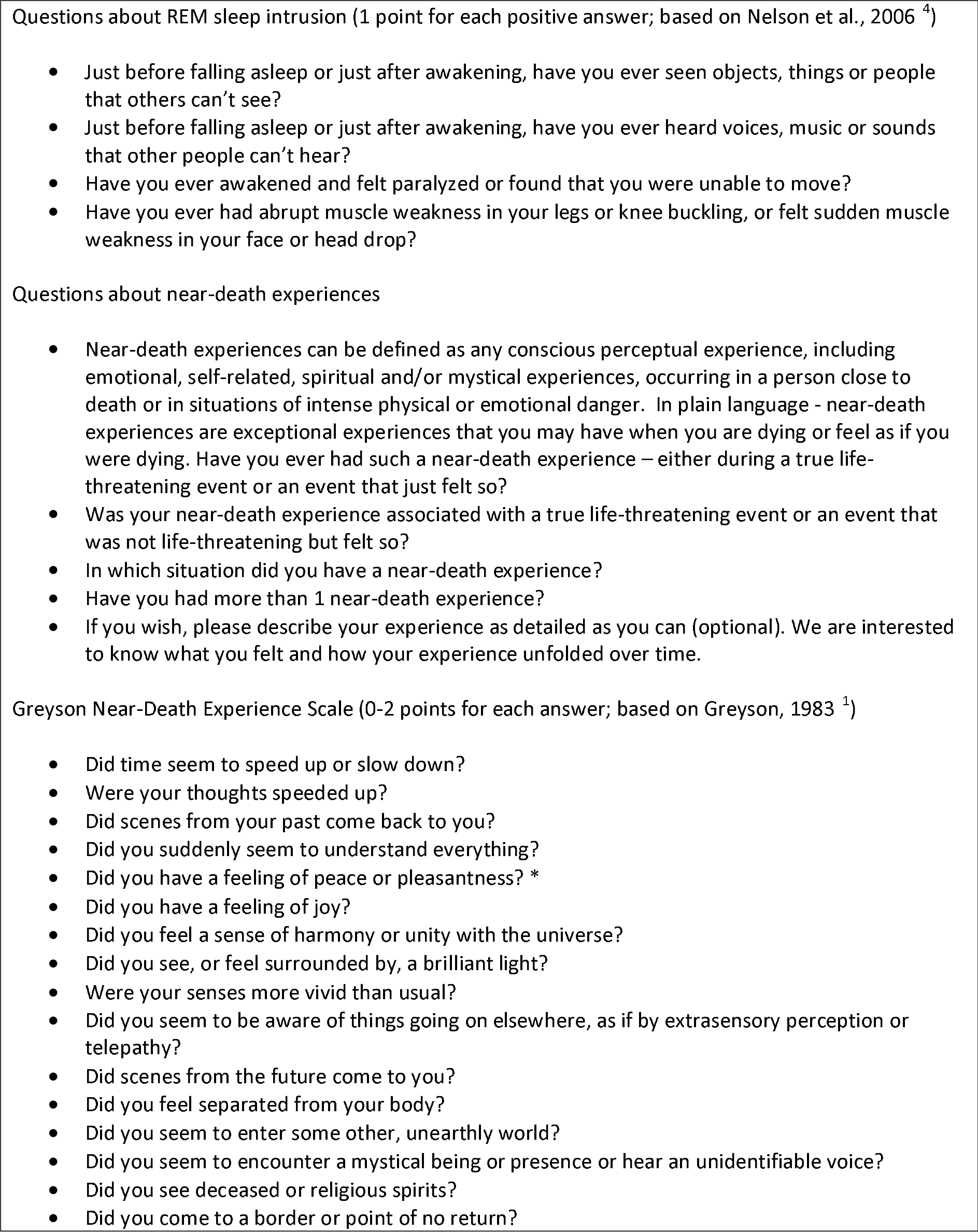
Questionnaire on REM sleep intrusion and near-death experiences. REM – rapid eye movements; * in contrast to the Near-Death Experience Scale, we also inquired about unpleasant experiences

## METHODS

### Hypotheses and research questions

The objectives of this study were two-fold.

- Primary objective: To estimate the frequency of near-death experiences and REM sleep intrusions reported in a large sample of adult humans collected from an online crowdsourcing service.
- Secondary objective: To test the hypothesis that people who report a near-death experience have a greater frequency of REM sleep intrusions.

### Study design

An online platform, Prolific Academic (https://prolific.ac/), was used to recruit a large global sample of lay people. Prolific Academic is a crowdsourcing online platform to recruit human subjects for research that compares favorably with Amazon’s Mechanical Turk in terms of honesty and diversity of participants and data quality. ^5,6^ Participants were recruited without any filters except for English language and age ≥18 years.

We asked participants to complete a questionnaire comprising demographic information on age, gender, employment status and place of residence, followed by 4 questions about REM sleep intrusion. Participants were then asked if they ever had had a near-death experience. If not, the survey ended here; if yes, we continued to inquire about this experience in detail, including all 16 items of the Greyson Near-Death Experience Scale (GNDES), the most widely used standardized tool to identify, confirm and characterize near-death experiences in research.^1^ In addition to multiple choice answers, participants were given the possibility to describe their experience in their own words. Prior to the start of the survey, participants were instructed that their monetary reward was fixed (see below), regardless of whether they would report having had a near-death experience or not. See **Table 1** for details.

### Statistics

We estimated the number of participants required to be 384, using a very high population size (300.000.000), a confidence level of 95% and a margin of error of 5%. However, as previous studies have estimated that 5-15% of the population have had a near-death experience,^7–9^ we decided to enroll ca. 1000 participants to identify an estimated 50-100 near-death experiences.

Data were analyzed using R (R 3.4.1, R Development Core Team [2008], Vienna, Austria). Categorical variables were analyzed using a chi-squared test. Continuous variables were compared using Student’s t-tests. We calculated odds ratios for having near-death experiences with or without co-occurrence of REM sleep intrusion and performed multivariate logistic regression analysis to correct for age, gender, employment status, place of residence, and whether the situation in which an experience was made was perceived as life-threatening or not. To adjust for multiple testing, the alpha level was set at 0.01.

To prevent HARKing (Hypothesizing After the Results are Known), ^10^ we pre-registered the study, including all objectives, with the Open Science Framework (https://osf.io/ykr3g).

### Ethics

Participants gave consent for publication of their (anonymous) data. Participation was anonymous, voluntary and restricted to those ≥18 years. Participants received a monetary reward upon completion of the survey, following the platform’s ethical rewards principle (≥ $6.50/h). The Ethics Committee of the Capital Region of Denmark waives approval for online surveys (Section 14 (1) of the Committee Act. 2; http://www.nvk.dk/english).

## RESULTS

We recruited 1034 lay people from 35 countries (mean age 32.7 years, SD 11.3 years; 59% female; 79% fully or part-time employed or in training), most of which were residing in Europe and North America. **Table 2** and **Figure 1** provide epidemiological information.

**Table 2.**
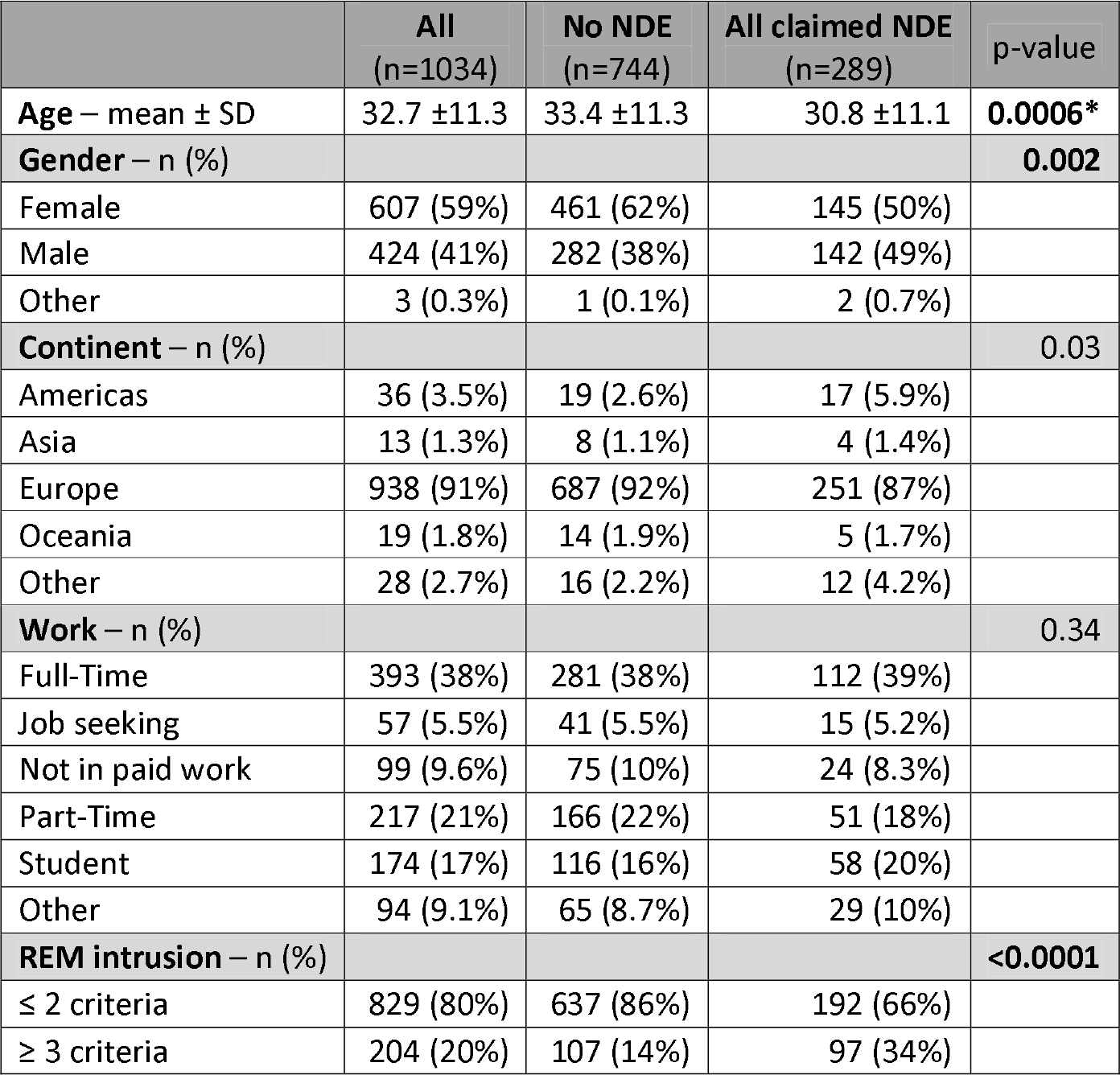
Demographic data and prevalence of REM sleep intrusion. To adjust for multiple testing, the alpha level was set to 0.01. Significant p values are shown in bold script. N – number of participants; NDE – near-death experiences; REM – rapid eye movements; SD – standard deviation; * when comparing “No NDE” (n=744) with confirmed near-death experiences with a Greyson NDE Scale score ≥ 7 (n=106; see Table 3), this significance is lost (p-value = 0.256).

**Figure 1.**
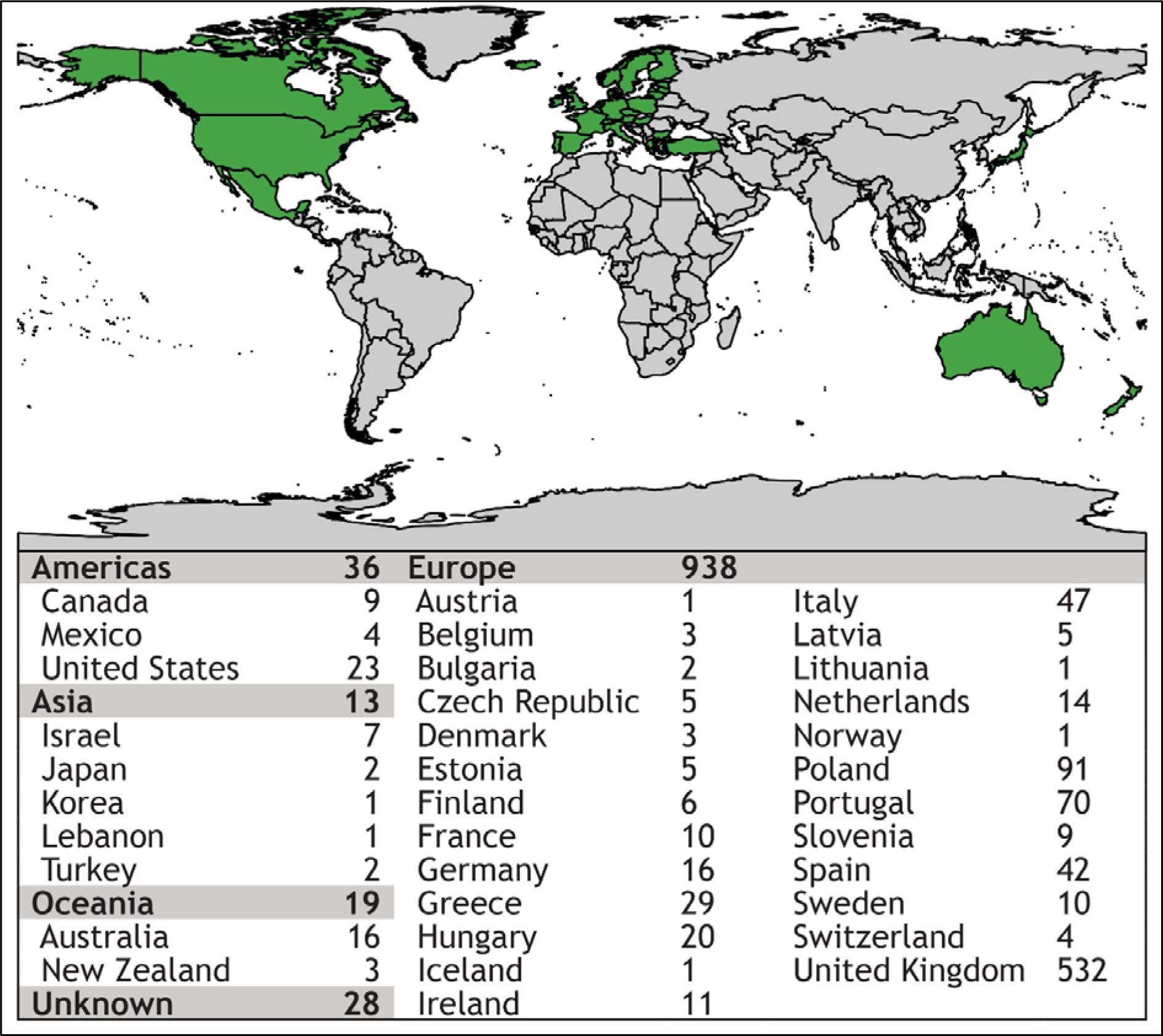
Map showing places of residency of survey participants. Using an online crowdsourcing platform, we recruited 1.034 lay people from 35 countries on 4 continents, the majority from Europe and North America

### Near-death experiences: Prevalence and semiology

Two-hundred eighty-nine participants (28%; CI 95% 25-31%) claimed to have had a near-death experience, 106 of which reached the threshold of 7 points or more on the GNDES (37%; CI 95% 31-43%). Thus, confirmed near-death experiences were reported by 106 of 1034 participants (10%; CI 95% 8.5-12%) (**Figure 2**).

**Figure 2.**
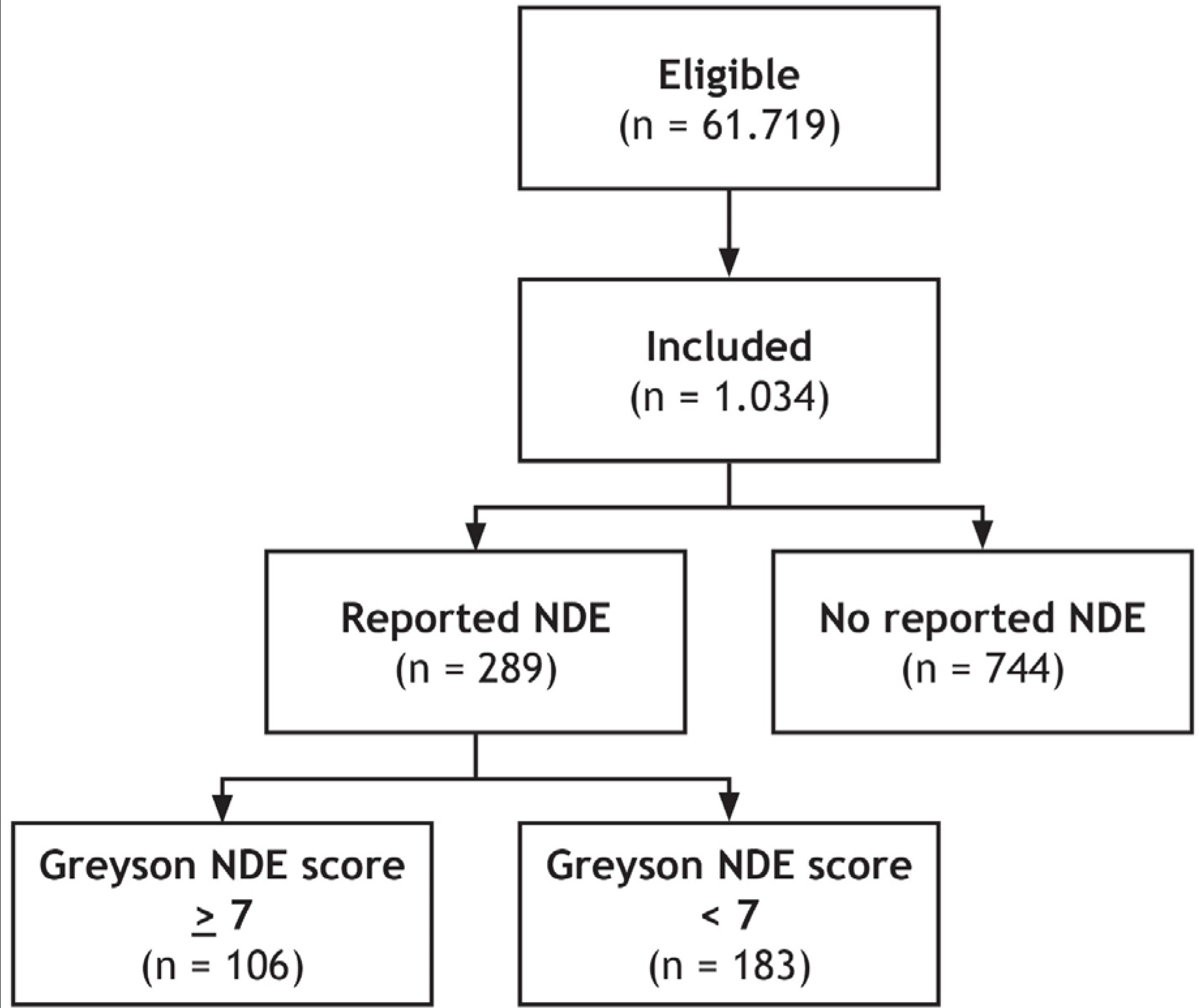
Schematic overview of study design. Of 61.719 eligible lay people registered with Prolific Academic (https://prolific.ac/; accessed on January 22, 2019), we enrolled 1.034 participants; 106 (10%; CI95% 8.5-12%) of whom reported a near-death experience that fulfilled established criteria (Greyson Near-Death Experience Scale score of 7 or higher). N = number of participants; NDE – near-death experience

Participants perceived the situation in which they had their experience slightly more often as truly life-threatening (n=158; 55%) than not truly life-threatening (n=131; 45%), and this was irrespective of whether their experience met criteria for a near-death experience according to the GNDES or not (p=0.55; **Table 3**).

**Table 3.** Participants claiming a near-death experience, analyzed according to Greyson Near-Death Experience Scale score. A score of ≥ 7 confirms the reported experience as a near-death experience. IQR – interquartile range; n – number of participants; NDE – near-death experience(s); REM – rapid eye movements; SD – standard deviation; significant p-values are shown in bold script; *excluding participants reporting that their experience was neither pleasant nor unpleasant

|  | **All claimed NDE**<br>(n=289) | **Greyson NDE score < 7**<br>(n=183) | **Greyson NDE score $\geq 7$**<br>(n=106) | p-value |
| --- | --- | --- | --- | --- |
| **Greyson NDE score – median (IQR)** | 5 (3;8) | 4 (3;5) | 9 (8;14) |  |
| Age – mean ± SD | 30.8 ±11.1 | 30.0 ±10.7 | 32.0 ±11.6 | 0.14 |
| **Gender – n (%)** |  |  |  | **0.34** |
| Female | 145 (50%) | 98 (54%) | 47 (44%) |  |
| Male | 142 (49%) | 84 (46%) | 58 (55%) |  |
| Other | 2 (0.7%) | 1 (0.5%) | 1 (0.9%) |  |
| **REM intrusion – n (%)** |  |  |  | **0.0003** |
| $\leq 2$ criteria | 192 (66%) | 136 (74%) | 56 (53%) | |
| $\geq 3$ criteria | 97 (34%) | 47 (26%) | 50 (47%) | |
| **Life-threatening event – n (%)** |  |  |  | 0.55 |
| Yes | 158 (55%) | 103 (56%) | 55 (52%) |  |
| No | 131 (45%) | 80 (44%) | 51 (48%) |  |
| **Feelings associated with NDE \* – n (%)** | (n=230) | (n=153) | (n=77) | **<0.0001** |
| Unpleasant | 168 (73%) | 132 (86%) | 36 (47%) |  |
| Pleasant | 62 (27%) | 21 (14%) | 41 (53%) |  |

Near-death experiences and experiences with ≤ 6 points on the GNDES occurred in the following situations, listed with decreasing frequency: motor accident 26% (n=77), near-drowning 19% (n=56); intense grief or anxiety 17.5% (n=51), substance abuse 11.5% (n=33); psychological distress without organic disease 9.5% (n=28); physical violence other than combat 8% (n=24), critical illness 8% (n=23); childbirth complication 8% (n=23); suicide attempt 7% (n=20); anesthesia/medical procedure 7% (n=20%); cardiac arrest/heart attack 5.5% (n=16); meditation or prayer 5% (n=15); anaphylactic reaction 5% (n=14); combat situation 4% (n=11); syncope 2% (n=5); epileptic seizure 1.5% (n=4); and other 19% (n=56).

The most often reported symptoms were abnormal time perception (faster or slower than normal; reported by 252 participants; 87%); exceptional speed of thoughts (n=189; 65%); exceptional vivid senses (n=182; 63%); and feeling separated from one’s body, including out-of-body experiences (n=152; 52%).

Experiences that qualified as a true near-death experience according to the GNDES were perceived much more often as pleasant (n=41; 53%) than experiences that did not (n=21; 14%; p<0.0001; neutral experience excluded; **Table 3**).

Around one third of participants reporting a near-death experience or near-death-like experience stated having had two or three such experiences (n=92, 32%); and some even claimed to have had more than three (n=10; 3.5%).

### Near-death experiences and evidence for REM sleep intrusion

Evidence for REM sleep intrusion was much more common in people with experiences above the cut-off point of the Greyson Near-Death Experience Scale (n=50/106; 47%) than in people with experiences below this threshold (n=47/183; 26%) or in those without any such experience (n=107/744; 14%; p=<0.0001; **Tables 2** and **3**). Following multivariate regression analysis to adjust for age, gender, place of origin, employment status and perceived danger, this association remained highly significant; i.e. people with REM sleep intrusion were more likely to exhibit near-death experiences than those without REM sleep abnormalities (odds ratio 2.85; CI 95% 1.68-4.88; p=0.0001; **Table 4**).

**Table 4.**
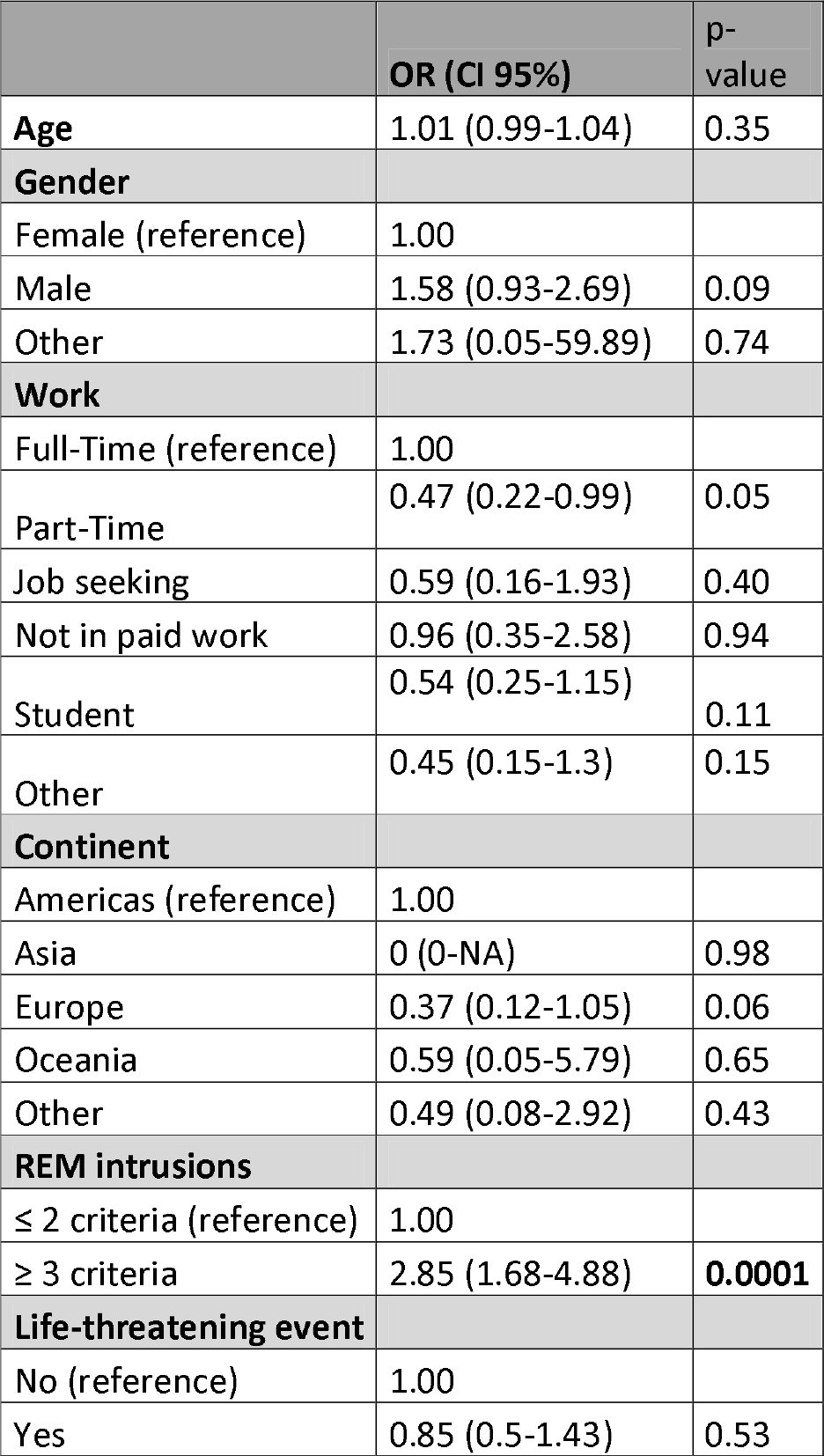
Multivariate logistic regression and odds ratios for having a near-death experience (Greyson Near-Death Experience Scale ≥ 7). To adjust for multiple testing, the alpha level was set to 0.01. CI – confidence interval; n – number of participants; OR – odds ratio; REM – rapid eye movements; significant p-values are shown in bold script

|  | **OR (CI 95%)** | **p-value** |
| --- | --- | --- |
| **Age** | 1.01 (0.99-1.04) | 0.35 |
| **Gender** |  |  |
| Female (reference) | 1.00 |  |
| Male | 1.58 (0.93-2.69) | 0.09 |
| Other | 1.73 (0.05-59.89) | 0.74 |
| **Work** |  |  |
| Full-Time (reference) | 1.00 |  |
| Part-Time | 0.47 (0.22-0.99) | 0.05 |
| Job seeking | 0.59 (0.16-1.93) | 0.40 |
| Not in paid work | 0.96 (0.35-2.58) | 0.94 |
| Student | 0.54 (0.25-1.15) | 0.11 |
| Other | 0.45 (0.15-1.3) | 0.15 |
| **Continent** |  |  |
| Americas (reference) | 1.00 |  |
| Asia | 0 (0-NA) | 0.98 |
| Europe | 0.37 (0.12-1.05) | 0.06 |
| Oceania | 0.59 (0.05-5.79) | 0.65 |
| Other | 0.49 (0.08-2.92) | 0.43 |
| **REM intrusions** |  |  |
| $\leq 2$ criteria (reference) | 1.00 | |
| $\geq 3$ criteria | 2.85 (1.68-4.88) | **0.0001** |
| **Life-threatening event** |  |  |
| No (reference) | 1.00 |  |
| Yes | 0.85 (0.5-1.43) | 0.53 |

Selected written reports from participants can be found in **Tables 5** and **6**. Raw data are provided in the *online supplemental files*.

**Table 5.**
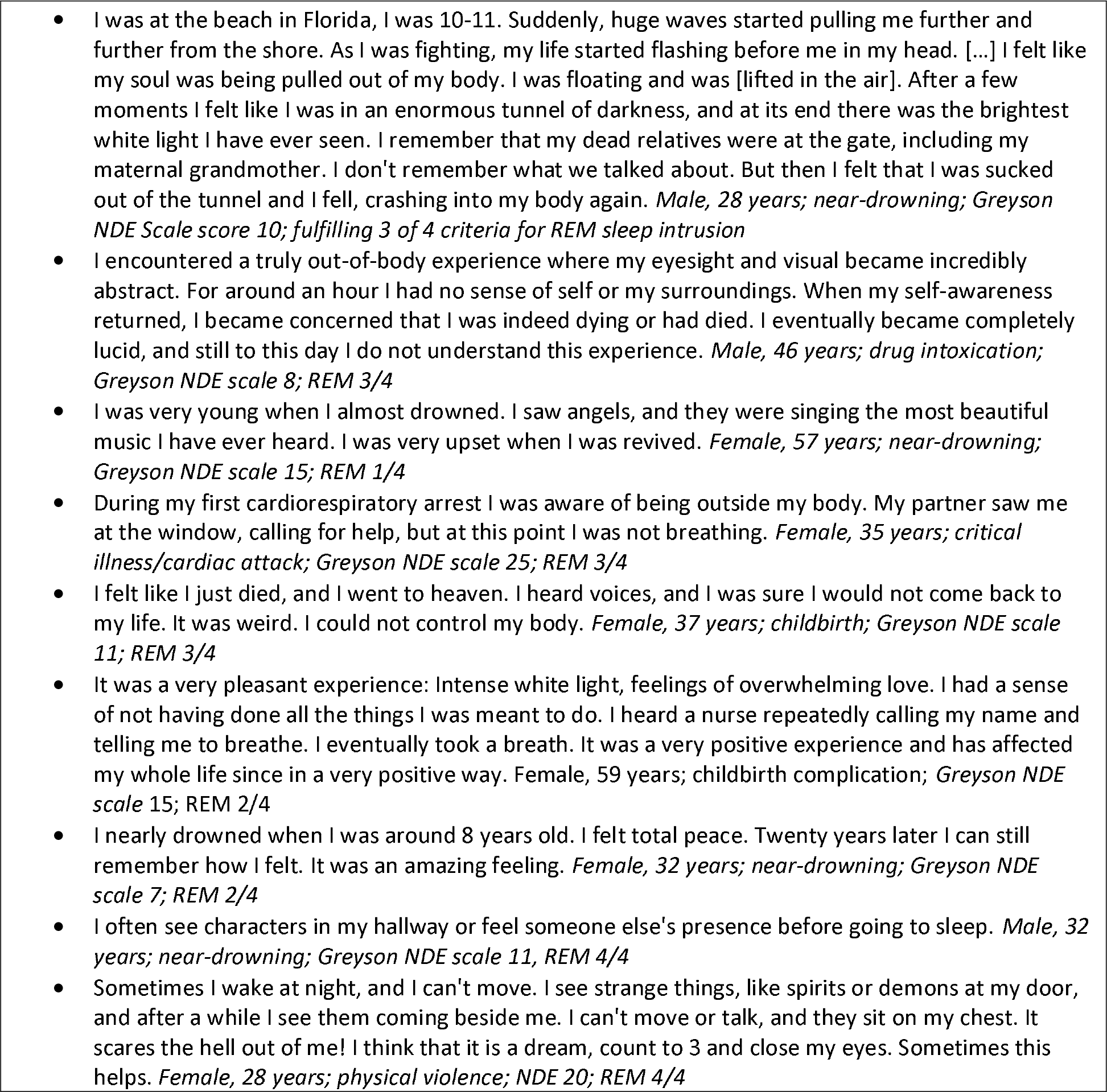
Selected reports from participants with an experience that reached the threshold of ≥7 points on the Greyson NDE scale to qualify as a near-death experience. Note that the last two comments describe instances that are highly suggestive of REM sleep disturbance, including visual hypnogogic hallucinations and sleep paralysis, rather than the near-death experience both participants reported to have had. Comments are edited for clarity and spelling.

**Table 6.**
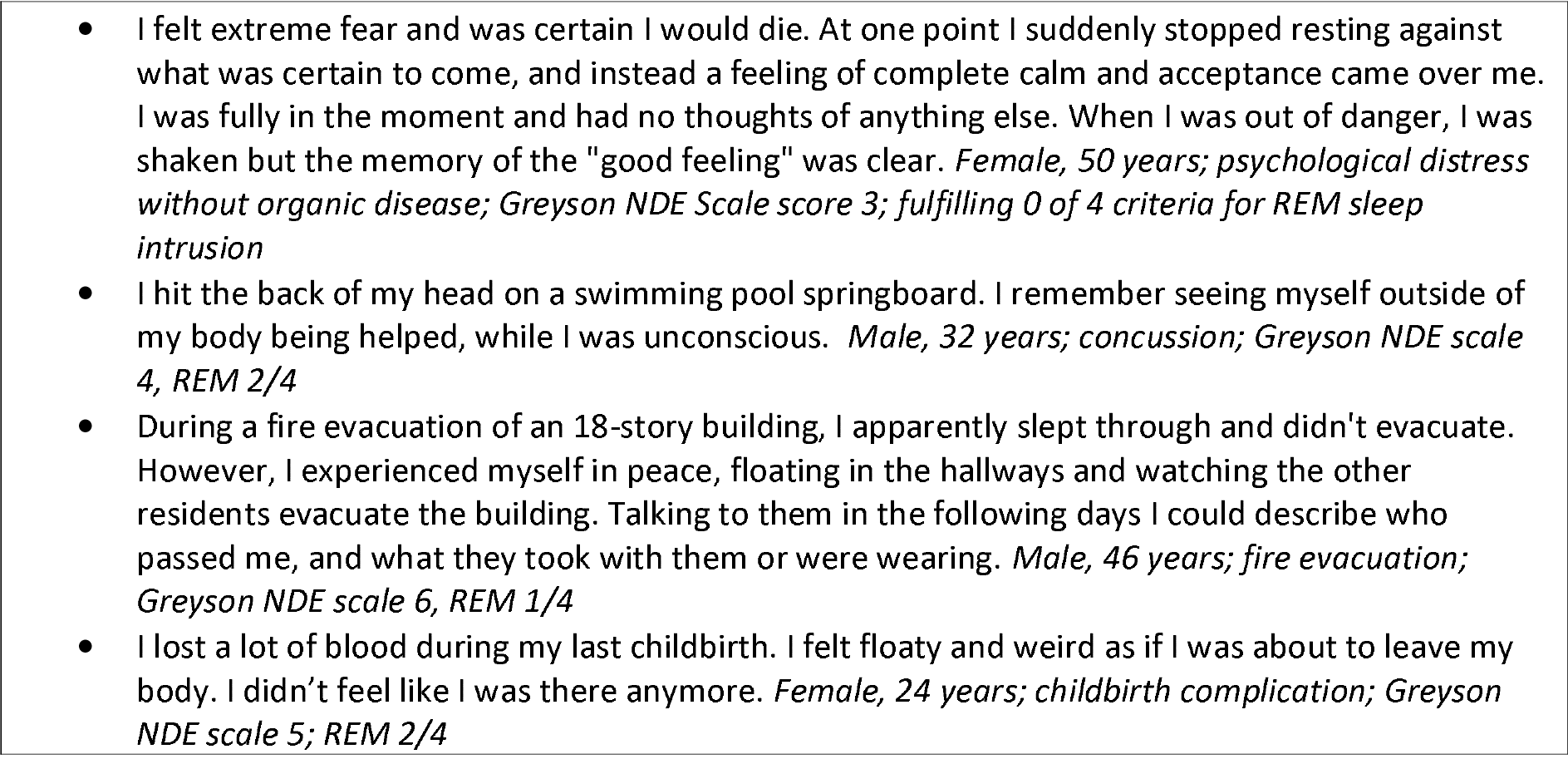
Selected reports from participants with an experience below the threshold of ≥7 points on the Greyson NDE scale. Comments are edited for clarity and spelling.

## DISCUSSION

Using crowdsourcing methods, we have shown that 1 out of 10 people from a large sample of 35 countries have had a confirmed near-death experience (10%; CI 95% 8.5-12%). This estimate is slightly higher than what was reported in three previous studies using traditional interview-based surveys in the US (5%)^7^, Australia (8%)^9^ and Germany (4%)^8^; of note, none of those studies validated reports with the GNDES. Experiences that did not fulfill criteria for a near-death experience were roughly twice as common.

Similar to previous reports, we found that near-death experiences occur in various cultures and nationalities and irrespective of employment status, age and gender.^2,3,11–13^ However, unlike previous reports in which near-death experiences were almost always associated with peacefulness and well-being,^2,3,11–13^ we found a much higher rate of people stating that their experience was indeed unpleasant. Although experiences with a cut-off score of at least 7 points on the GNDES were more often pleasant (53%) than experiences with a lower score (14%; p<0.0001), almost half of all near-death experiences were labelled as stressful. We believe this is a much more realistic picture. The discrepancy between the present and previous studies is likely due to methodological limitations of the GNDES that addresses only pleasant feelings but does not, in contrast to our questionnaire, record negative emotions.^1^

Also unlike previous studies,^2,3,11–13^ we found that near-death experiences occurred equally likely in truly life-threatening situations and situations that only just felt so. Again, we think this reflects a more representative picture, since our study, in contrast to most others, draws inferences from a large cross-sectional sample of unprimed lay people and not from retrospective or prospective observations in specific populations such as cardiac arrest survivors.^3,14^ Still, these data substantiate previous reasoning^12^ that near-death experiences are real experiences and not merely products of fantasy proneness: People with confirmed near-death experiences did not perceive their situations as more dangerous than those without such experiences which argues against tendencies towards overdramatizing.

The most important finding, however, is the association of near-death experiences with REM sleep intrusion. Following multivariate analysis, REM sleep intrusion was the only factor that remained significantly correlated with near-death experiences (and indeed very much so: p=0.0001). This finding corroborates and extends data from a previous case-control study, in which Nelson and co-workers assessed the life-time prevalence of REM sleep intrusion in 55 humans with near-death experiences compared with that in age- and sex-matched controls. Sleep-related visual and auditory hallucinations and sleep paralysis assessed by a questionnaire akin to that used in our study were substantially more common in cases with a near-death experience. The authors concluded that under circumstances of peril, near-death experiences are more likely in people with a tendency towards REM sleep intrusion and that REM sleep intrusion might explain much, if not all, of the semiology of these experiences. Indeed, as shown in **Table 5**, two participants from our study gave spontaneous reports of classic REM sleep disturbances (rather than reporting their near-death experience as requested) akin to those seen in people with narcolepsy.^15,16^

From a biological perspective, REM sleep intrusion as an explanation for near-death experiences appears to make sense. Recent observations into the physiology of the dying human brain are very noteworthy in this regard.^17,18^ Dreier and co-workers from the COSBID group (Co-Operative Studies on Brain Injury Depolarizations) performed recordings with subdural electrode strips or intraparenchymal electrode arrays in 9 patients with devastating brain injury and a Do Not Resuscitate-Comfort Care order. Following terminal extubation, the authors noted a decline in brain tissue partial pressure of oxygen and circulatory arrest. Of note, silencing of spontaneous cortical electrical activity developed simultaneously across regional electrode arrays in 8 patients. As suggested by the authors, “isoelectricity of brain activity develops as neurons hyperpolarize to reduce energy consumption as a final survival strategy”.^18^ This silencing, which the authors term nonspreading depression, was trailed by terminal spreading depolarizations starting to propagate between electrodes 2 to 6 minutes after onset of the final drop in cerebral perfusion and between 15 seconds and 4.5 minutes after nonspreading depression.^18^

Metaphysical speculations omitted, this state of cortical dying seems incompatible with preserved function of hippocampi and large-scale memory networks that is mandatory for the formation and storage of elaborate memories such as near-death experiences, let alone recovery of consciousness and cognition to report these events. In line with this reasoning, near-death experiences in our study were almost equally distributed between true life-threatening situations and situations that just felt so and in which brain injury was very unlikely to occur. We thus suggest that near-death experiences are not fully terminal or pre-terminal events but rather reflect physiological brain states with functionally and structurally preserved neuronal networks and sudden onset of REM sleep-like features.

An online study such as this one has limitations that should be acknowledged. ^5,6^ First, complex clinical and ethical notions are impossible to fully implement in a survey form. Second, although we assessed various demographic factors, there are many with potential importance such as religiosity that we did not assess. (Although we did assess religiosity in a previous online survey using the same crowdsourcing platform and found that most participants have a secular background.^19^ Also, of note, religiosity appears to be of little importance to near-death experiences.^20^)

On the positive side, the anonymous nature of an online survey limits the influence of psychological bias; there is no incentive to please the investigator by inventing or exaggerating memories. (There was no monetary incentive either, as participants were instructed that their reimbursement was the same irrespective of whether they would report a near-death experience or not). Further, this is the first systematic study on the prevalence of near-death experiences and REM sleep intrusion in an unselected sample of adult lay people. We recruited a much larger sample size than what typically can be achieved with lab-based behavioral testing or mail-based questionnaires. Although we were unable to recruit participants from Africa and Asian participants were underrepresented, this was a truly global sample with respondents from more than 35 countries on 4 continents, which strengthens the validity and generalizability of our results.

## CONCLUSIONS

The prevalence of near-death experiences in the public is around 10%. Whereas age, gender, place of residence, employment status and factual danger of the situation do not appear to influence the frequency with which near-death experiences occur, there is a significant association with REM sleep intrusion. This observation fits well with the notion that despite imminent threat to life, brain physiology must be well-preserved to perceive these fascinating experiences and store them as long-term memories.

**Author contribution:**
DK - study concept, acquisition of data, analysis and interpretation, writing of the manuscript, critical revision for important intellectual content, and approval of final manuscript. MHO - analysis and interpretation, critical revision for important intellectual content, and approval of final manuscript.

